# Bayesian estimation of Lassa virus epidemiological parameters: implications for spillover prevention using wildlife vaccination

**DOI:** 10.1101/841403

**Authors:** Scott L. Nuismer, Christopher H. Remien, Andrew Basinski, Tanner Varrelman, Nathan Layman, Kyle Rosenke, Brian Bird, Michael Jarvis, Peter Barry, Elisabeth Fichet-Calvet

## Abstract

Lassa virus is a significant burden on human health throughout its endemic region in West Africa, with most human infections the result of spillover from the primary rodent reservoir of the virus, the natal multimammate mouse, *M. natalensis*. Here we develop a Bayesian methodology for estimating epidemiological parameters of Lassa virus within its rodent reservoir and for generating probabilistic predictions for the efficacy of rodent vaccination programs. Our approach uses Approximate Bayesian Computation (ABC) to integrate mechanistic mathematical models, remotely-sensed precipitation data, and Lassa virus surveillance data from rodent populations. Using simulated data, we show that our method accurately estimates key model parameters, even when surveillance data are available from only a relatively small number of points in space and time. Applying our method to previously published data from two villages in Guinea estimates the time-averaged *R*_0_ of Lassa virus to be 1.658 and 1.453 for rodent populations in the villages of Bantou and Tanganya, respectively. Using the posterior distribution for model parameters derived from these Guinean populations, we evaluate the likely efficacy of vaccination programs relying on distribution of vaccine-laced baits. Our results demonstrate that effective and durable reductions in the risk of Lassa virus spillover into the human population will require repeated distribution of large quantities of vaccine.

**Author Summary:** Lassa virus is a chronic source of illness throughout West Africa, and is considered to be a threat for widespread emergence. Because most human infections result from contact with infected rodents, interventions that reduce the number of rodents infected with Lassa virus represent promising opportunities for reducing the public health burden of this disease. Evaluating how well alternative interventions are likely to perform is complicated by our relatively poor understanding of viral epidemiology within the reservoir population. Here we develop a novel statistical approach that couples mathematical models and viral surveillance data from rodent populations to robustly estimate key epidemiological parameters. Applying our method to existing data from Guinea yields well-resolved parameter estimates and allows us to simulate a variety of rodent vaccination programs. Together, our results demonstrate that rodent vaccination alone is unlikely to be an effective tool for reducing that public health burden of Lassa fever within West Africa.

## Introduction

Lassa virus is a zoonotic pathogen endemic to West Africa where it poses a significant burden on human health (Buckley et al. 1970). Although only a relatively small proportion of human cases result in severe symptoms and mortality, human infection is common, with 8-52% of the human population within Sierra Leone seropositive (McCormick et al. 1987; Bausch et al. 2001; Dan-Nwafor et al. 2019), indicative of past infection by Lassa virus. In addition to being a chronic source of illness throughout West Africa, Lassa virus is considered to be a threat for widespread emergence and is recognized by the World Health Organization as a priority pathogen (Sweileh 2017).

Human infection with Lassa virus occurs primarily through contact with excretions of the primary reservoir species, the natal multimammate mouse, *M. natalensis* (McCormick et al. 1987). Within Lassa endemic regions of West Africa, *M. natalensis* frequently inhabit human dwellings and trapping studies have demonstrated that up to 30% of *M. natalensis* individuals can be PCR positive for Lassa virus (Fichet-Calvet et al. 2007; Safronetz et al. 2013). Recent modeling and genetic studies have confirmed the primary importance of zoonotic transmission, providing evidence that human to human transmission is rare outside of hospital settings (Lo Iacono et al. 2015; Siddle et al. 2018; Kafetzopoulou et al. 2019). Because most human infections result from contact with infected rodents, strategies for reducing human infection have focused on reducing human contact with the reservoir species, *M. natalensis*, reducing the reservoir population as a whole through trapping or poisoning, or reducing the proportion of infected rodents through vaccination (Mendoza et al. 2018; Saez et al. 2018; Marien et al. 2019).

Recent studies investigating the efficacy of rodent removal using annual application of a rodenticide in Guinea demonstrated poisoning could yield substantial transient reductions in the density of *M. natalensis* (Saez et al. 2018). Specifically, this study applied rodenticide during the dry season over a four-year period and evaluated trapping success at the beginning and end of each application period. Although trapping success (and presumably rodent density) declined by the end of each application period, populations rapidly rebounded to their pre-treatment levels in all but the fourth year. Thus, the extent to which rodenticide application can yield durable reductions in rodent density remains unclear. In contrast to rodent removal, no experimental studies examining the impact of rodent vaccination exist because vaccines targeting Lassa virus in the reservoir population are under development, but not yet available. As a consequence, efforts to predict how well vaccination campaigns might work have relied on computer simulations (e.g., Marien et al. 2019). Using individual based simulations to predict the effectiveness of culling and vaccination, Marien et al. (2019) found that Lassa virus could be eliminated from its reservoir population only through continuous rodent control or vaccination. Because these conclusions rest on informal model parameterization, do not integrate seasonal variation in reproduction, and investigate only a small range of possible vaccination strategies, their generality remains unclear.

Here we develop a robust Bayesian methodology for estimating key epidemiological parameters of Lassa virus within its natural reservoir, the natal multimammate mouse, *Mastomys natalensis,* that couples mechanistic mathematical models, remotely sensed precipitation data, and rodent capture data from two villages in Guinea using Approximate Bayesian Computation (ABC). We use this Bayesian approach to estimate individual model parameters as well as the time averaged *R*_0_ for Lassa virus within two villages in Guinea for which time-series data is available (Fichet-Calvet et al. 2007; Fichet-Calvet et al. 2014). The time-averaged value of *R*_0_ measures the average number of new Lassa virus infections produced by an infected rodent if it were introduced into an entirely susceptible population over the 17-month period of data collection. Repeated sampling from the posterior distribution allowed us to generate probabilistic predictions for the efficacy of vaccination campaigns and to identify the optimal timing of vaccine distribution.

## Methods

### Mathematical Model

We model the coupled ecological and epidemiological dynamics of *M. natalensis* and Lassa virus in a metapopulation consisting of a set of 𝒫 populations connected by migration. We assume the age structure of the reservoir population, *M. natalensis*, can be well-described by three discrete life stages: pup, pre-reproductive juvenile, and reproductive adult. The epidemiological dynamics of Lassa virus are assumed to follow an SIR model where individual rodents are either susceptible to Lassa infection (S), currently infected by Lassa virus and infectious to other rodents (I), or recovered from Lassa virus infection and immune to further infection (R). We do not include the complication of modeling an exposed class (e.g., Marien et al. 2019) because experimental studies suggest viral shedding may commence rapidly (2-3 days) after exposure (Rosenke, K. Unpublished data). Our model allows for horizontal transmission of Lassa virus among classes as well as vertical transmission from mother to offspring and transmission of protective maternal antibodies. We include the possibility of vertical transmission and maternal antibody transfer because both have been demonstrated in related arenaviruses and included in previous models exploring the efficacy of Lassa control measures (Marien et al. 2019). Within a location, x, we assume individuals encounter one another at random and that juveniles and adults move between different locations *x* and *y* at per capita rates *m*_*J*,*x*,*y*_ and *m*_*A*,*x*,*y*_, respectively. Both density-dependent and density-independent mortality are included as both have been demonstrated to be important in *M. natalensis* population dynamics (Leirs et al. 1997). Together, these assumptions lead to the following system of differential equations describing the dynamics of Lassa virus infection within a single geographic location, x:

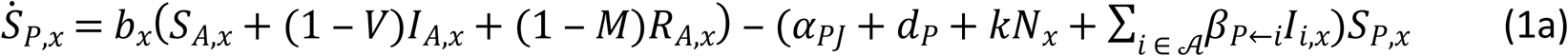

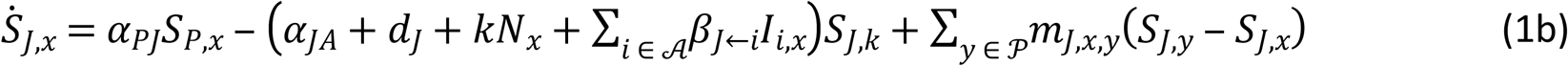

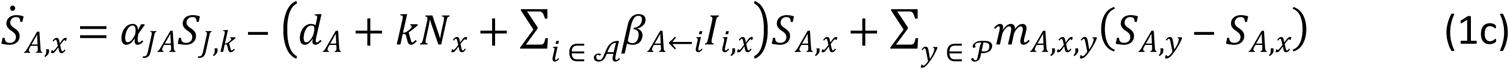

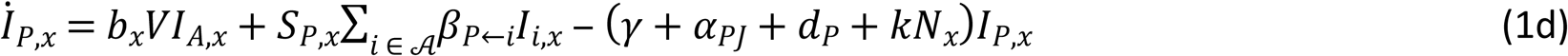

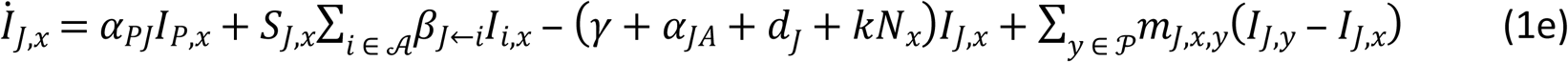

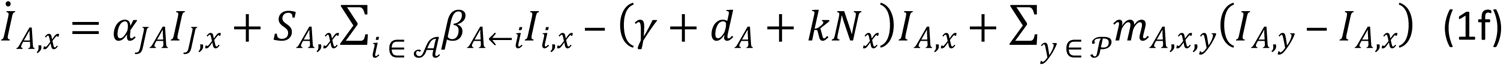

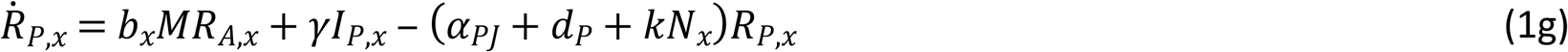

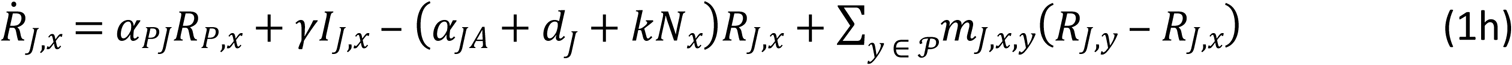

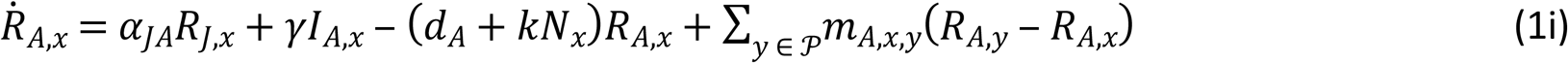

where all parameter and variables are described in Table 1.

**Table 1.** Model variables and parameters

| Variables/Parameters | Biological interpretation |
| --- | --- |
| $\mathcal{A}$ | The set of all age classes |
| $\mathcal{P}$ | The set of all populations |
| $N_x$ | The total number/density of *M. natalensis* juveniles and adults in location $x$ |
| $S_{i,x}$ | The number/density of Lassa virus susceptible *M. natalensis* in age class $i$ at location $x$ (PCR-/Sero-) |
| $I_{i,x}$ | The number/density of Lassa virus infected *M. natalensis* in age class $i$ at location $x$ (PCR+/Sero- and PCR+/Sero+) |
| $R_{i,x}$ | The number/density of Lassa virus recovered/resistant *M. natalensis* in age class $i$ and location $x$ (PCR-/Sero+) |
| $b_x(t)$ | *M. natalensis* birth rate at location $x$ and time $t$ |
| $b_M$ | Maximum possible per capita birth rate for *M. natalensis* |
| $\rho$ | The sensitivity of *M. natalensis* birth rate to precipitation |
| $\omega$ | The number of days over which precipitation is averaged |
| $\bar{P}_x$ | Average precipitation at location $x$ over the preceding $\omega$ days |
| $k_x$ | Density dependent death rate of *M. natalensis* at location $x$ |
| $d_i$ | Density independent death rate of *M. natalensis* in age class $i$ |
| $\alpha_{ij}$ | Maturation rate from age class $i$ to age class $j$ |
| $\beta_{i \leftarrow j}$ | Transmission rate of Lassa virus from individuals in age class $j$ to individuals in age class $i$. Horizontal transmission is assumed to occur only among juveniles and adults. |
| $V$ | Probability of vertical transmission of Lassa virus infection |
| $M$ | Probability of maternal antibody transfer |
| $\gamma$ | Recovery rate from Lassa virus infection |
| $m_{i,x,y}$ | Rate of movement between populations x and y for age class $i$ |
| $\tau_i$ | Rate at which individuals in age class $i$ are trapped |

Because seasonal patterns of rainfall are known to influence reproduction in at least some *M. natalensis* populations (e.g., Leirs et al. 1997), our model allows the birth rate to fluctuate in response to precipitation. Specifically, we assume that the current birth rate of a population at location x depends on the average precipitation that has fallen at the location over the previous *⍵* days, *P̅*_*x*_. Allowing the sensitivity of current birth rate to average precipitation to be tuned by the parameter *ρ* yields the following function describing the birth rate at location x and time t:

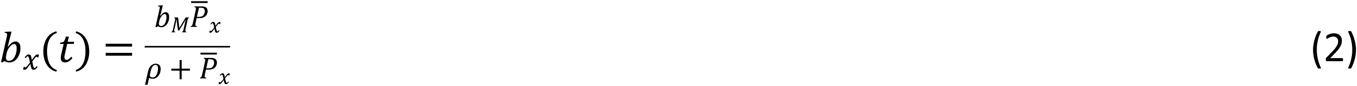

where *P̅*_*x*_ for each location x is calculated by averaging daily precipitation values provided by the CHIRPS 2.0 database (Funk et al. 2015) for the 0.05 degree grid square in which the latitude and longitude of the location x fall.

### Rodent Capture Data

We focus our inference procedure on data collected from a multi-year rodent trapping study conducted in Guinea between 2002 and 2005 (Fichet-Calvet et al. 2007; Fichet-Calvet et al. 2014). Rodents were trapped in two villages (Bantou and Tanganya) at multiple time points within each year using well established procedures. Each captured rodent was identified, aged using eye lens weight, sexed, and screened for LASV infection using both PCR and Serology with details as previously described (Fichet-Calvet et al. 2007; Fichet-Calvet et al. 2014). We translated the original data into our SIR modeling framework by classifying individuals as juvenile if age estimated from eye lens weight was less than 131.74 days and adult if estimated age was equal or greater than 131.74 days. We chose 131.74 days as the threshold because this is the average age of first reproduction for female *M. natalensis* within a captive colony of *M. natalensis* originally captured in Mali and housed at Rocky Mountain Laboratories (Rosenke, K. Unpublished data). No pups (pre-weaning life stage) were captured so data on this age class are unavailable. Individuals were classified as susceptible (S) if they were PCR and serologically negative for Lassa virus, infected (I) if they were PCR positive for Lassa virus, and recovered/resistant if they were PCR negative for Lassa virus but serologically positive. Using these definitions to translate the data into our modeling framework results in a time series for each village describing the number of juvenile and adult *M. natalensis* that are susceptible, infectious, or recovered at each sampling time point (Figure 1).

**Figure 1.**
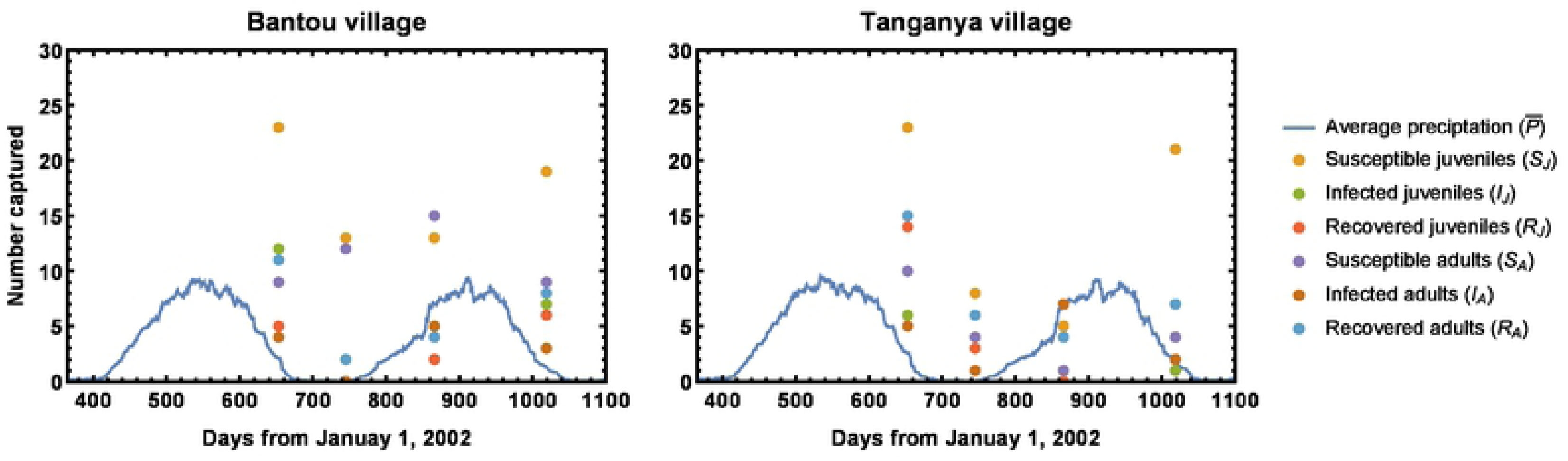
Time series data for the number of *M. natalensis* individuals captured within two villages in Guinea over five trapping sessions occurring between 2002-2005 (colored dots). The blue lines indicate the 60-day average precipitation values for each village. *M. natalensis* capture data comes from the study described in (Fichet-Calvet et al. 2007; Fichet-Calvet et al. 2014)and precipitation data comes from the CHIRPS 2.0 database. Although the original rodent trapping study included two additional dates, these were left out of our analyses because these dates did not have complete data on all classes or had substantially reduced trapping effort.

### Approximate Bayesian Computation

Because our model is too complex to solve analytically and does not allow expressions for the likelihood of observed time-series data to be explicitly formulated, we developed an Approximate Bayesian approach. Specifically, our mechanistic model is parameterized using Approximate Bayesian Computation (ABC). In brief, ABC works by: 1) drawing parameters at random from prior distributions informed by existing data, 2) simulating data under the model (i.e., a time series of *M. natalensis* abundance and infection status), and 3) placing the randomly drawn parameters into the posterior distribution only if the simulated data is sufficiently close to the real data. Repeating this process ultimately results in an approximate posterior distribution for model parameters (Beaumont 2010; Bertorelle et al. 2010; Csillery et al. 2010)

Our implementation of ABC relies on simulating the model (1-2) using parameters drawn from the prior distributions shown in Table 2 and daily precipitation data for each geographic location downloaded from the CHIRPS 2.0 database (Funk et al. 2015). We used the Gillespie algorithm (Gillespie 1977) to stochastically simulate the analogous continuous-time Markov chain version of equations (1-2), with simulations initiated 288 days prior to the start of rodent sampling to allow ecological and epidemiological dynamics to burn into a stationary distribution prior to comparing simulated and real data. To compare simulated values of rodent abundances to real data on numbers of rodents captured using traps, we implemented simulated rodent trapping experiments at each time point for which data was available. These simulated rodent capture experiments proceeded by drawing a random number from a binomial distribution for each rodent age/infectious class with the number of trials set to the number of trap nights in the trapping session and the probability of capture, *p*, set to the value:

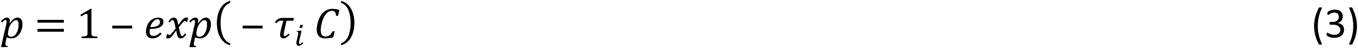

where *τ_i_* is the rate at which rodents in age class *i* are trapped and *C* is the simulated population density of the class. Thus, the greater the population density of a particular class, the more likely individuals of this class are to be captured by a trap over the course of an individual trap night.

**Table 2.**
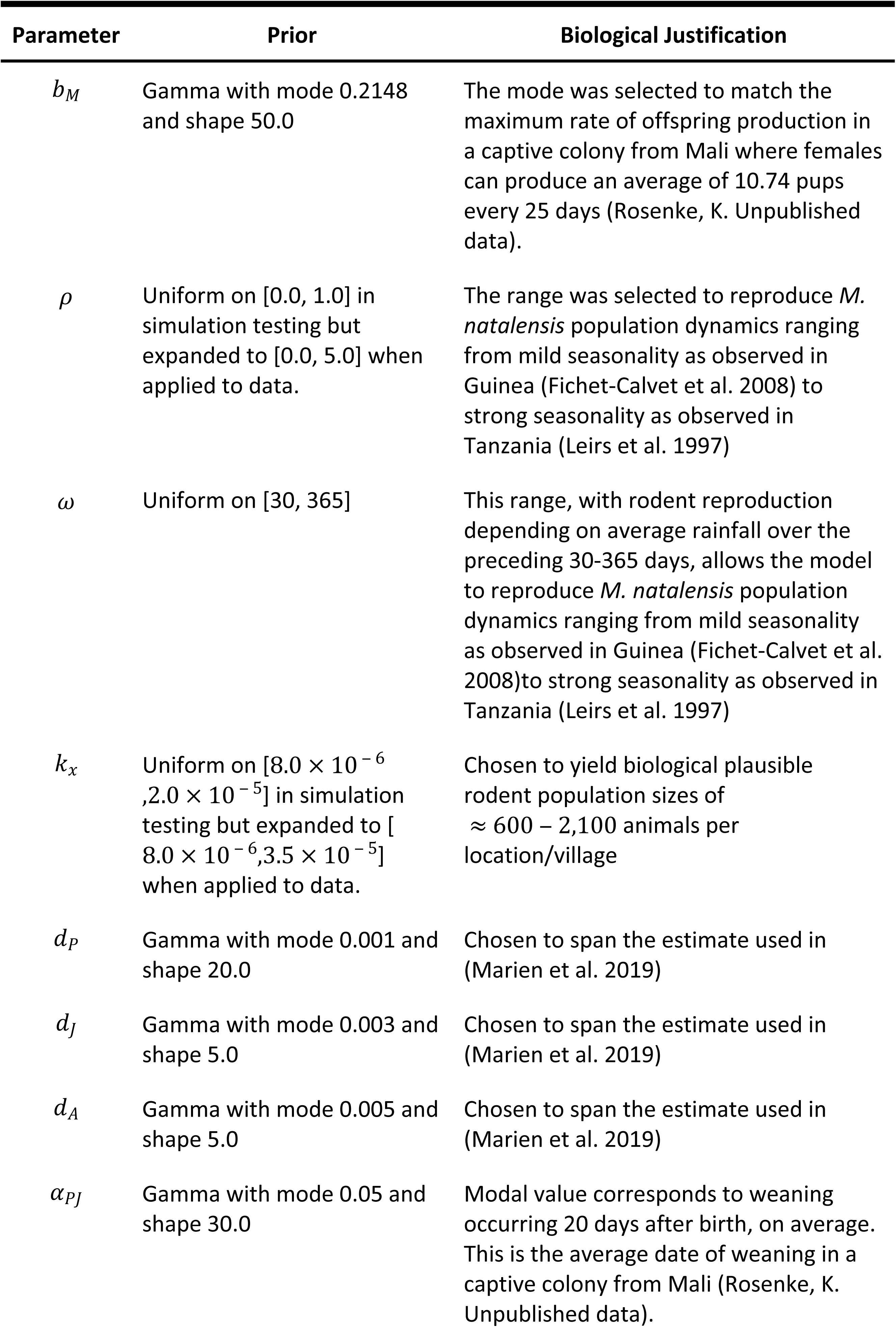
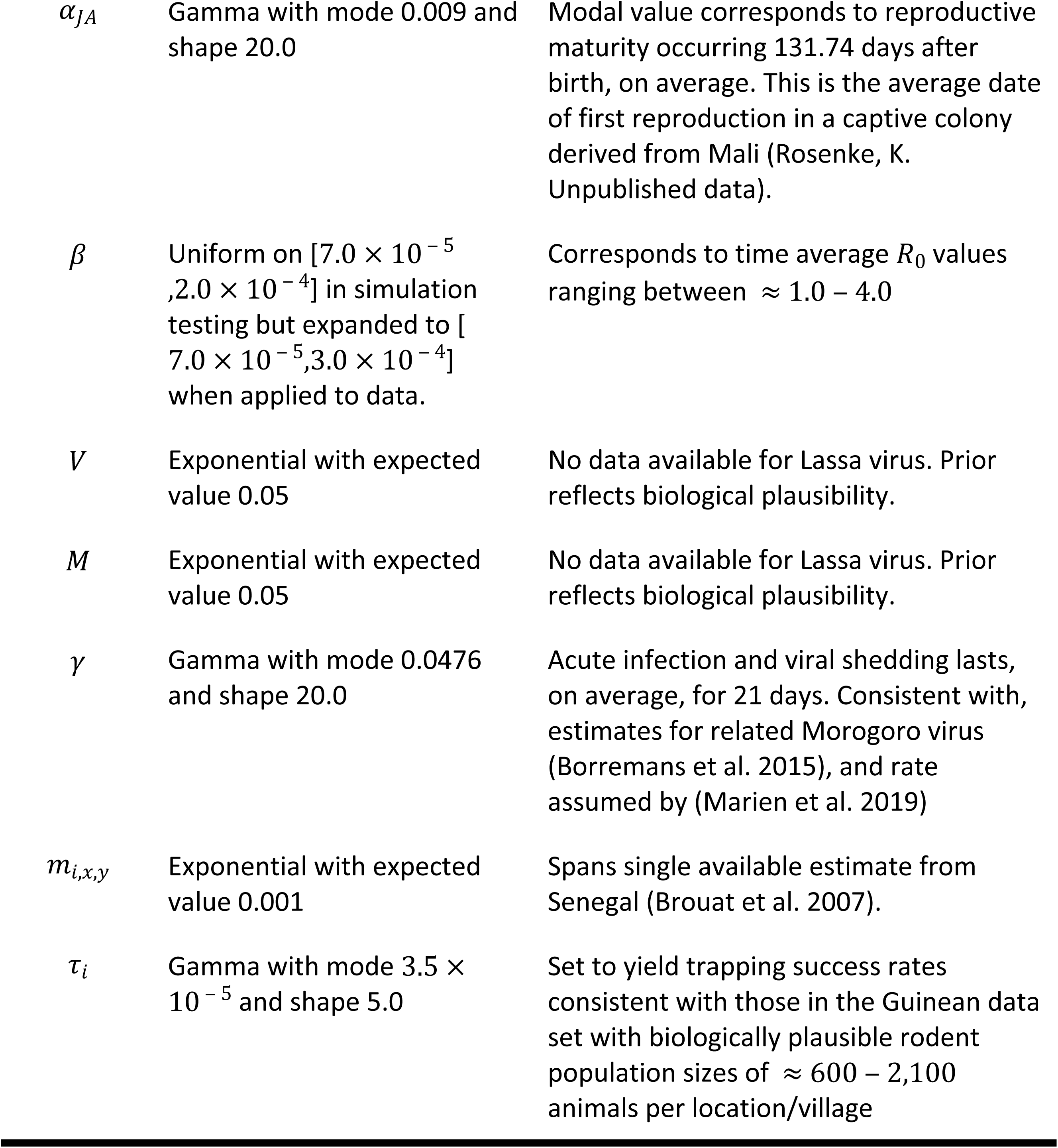
Prior distributions for model parameters

| Parameter | Prior | Biological Justification |
| --- | --- | --- |
| $b_M$ | Gamma with mode 0.2148 and shape 50.0 | The mode was selected to match the maximum rate of offspring production in a captive colony from Mali where females can produce an average of 10.74 pups every 25 days (Rosenke, K. Unpublished data). |
| $\rho$ | Uniform on [0.0, 1.0] in simulation testing but expanded to [0.0, 5.0] when applied to data. | The range was selected to reproduce *M. natalensis* population dynamics ranging from mild seasonality as observed in Guinea (Fichet-Calvet et al. 2008) to strong seasonality as observed in Tanzania (Leirs et al. 1997) |
| $\omega$ | Uniform on [30, 365] | This range, with rodent reproduction depending on average rainfall over the preceding 30-365 days, allows the model to reproduce *M. natalensis* population dynamics ranging from mild seasonality as observed in Guinea (Fichet-Calvet et al. 2008) to strong seasonality as observed in Tanzania (Leirs et al. 1997) |
| $k_x$ | Uniform on [$8.0 \times 10^{-6}$, $2.0 \times 10^{-5}$] in simulation testing but expanded to [$8.0 \times 10^{-6}$, $3.5 \times 10^{-5}$] when applied to data. | Chosen to yield biological plausible rodent population sizes of $\approx 600 - 2,100$ animals per location/village |
| $d_P$ | Gamma with mode 0.001 and shape 20.0 | Chosen to span the estimate used in (Marien et al. 2019) |
| $d_J$ | Gamma with mode 0.003 and shape 5.0 | Chosen to span the estimate used in (Marien et al. 2019) |
| $d_A$ | Gamma with mode 0.005 and shape 5.0 | Chosen to span the estimate used in (Marien et al. 2019) |
| $\alpha_{PJ}$ | Gamma with mode 0.05 and shape 30.0 | Modal value corresponds to weaning occurring 20 days after birth, on average. This is the average date of weaning in a captive colony from Mali (Rosenke, K. Unpublished data). |
| $\alpha_{JA}$ | Gamma with mode 0.009 and shape 20.0 | Modal value corresponds to reproductive maturity occurring 131.74 days after birth, on average. This is the average date of first reproduction in a captive colony derived from Mali (Rosenke, K. Unpublished data). |
| $\beta$ | Uniform on $[7.0 \times 10^{-5}, 2.0 \times 10^{-4}]$ in simulation testing but expanded to $[7.0 \times 10^{-5}, 3.0 \times 10^{-4}]$ when applied to data. | Corresponds to time average $R_0$ values ranging between $\approx 1.0 - 4.0$ |
| $V$ | Exponential with expected value 0.05 | No data available for Lassa virus. Prior reflects biological plausibility. |
| $M$ | Exponential with expected value 0.05 | No data available for Lassa virus. Prior reflects biological plausibility. |
| $\gamma$ | Gamma with mode 0.0476 and shape 20.0 | Acute infection and viral shedding lasts, on average, for 21 days. Consistent with, estimates for related Morogoro virus (Borremans et al. 2015), and rate assumed by (Marien et al. 2019) |
| $m_{i,x,y}$ | Exponential with expected value 0.001 | Spans single available estimate from Senegal (Brouat et al. 2007). |
| $\tau_i$ | Gamma with mode $3.5 \times 10^{-5}$ and shape 5.0 | Set to yield trapping success rates consistent with those in the Guinean data set with biologically plausible rodent population sizes of $\approx 600 - 2,100$ animals per location/village |

For each simulation, simulated trapping data and real trapping data were compared, and parameter combinations yielding simulated trapping data sufficiently close to the real trapping data were included in the posterior distribution. Specifically, parameter combinations were added to the posterior if two criteria were met. First, the total number of juvenile and adult animals captured in the simulated data must be within 25% of its true value, on average, across all trapping sessions and locations. Second, the sum of the absolute distance between frequencies of captured individuals belonging to infected juvenile (*I_J_*), infected adult (*I_A_*), juvenile recovered (*R_J_*), and adult recovered (*R_A_*) classes in the simulated and real data must be within 0.5 of its true value, on average, across all trapping sessions and locations. Our analyses of simulated data demonstrated that five individual model parameters can be reliably estimated: 1) the rate of horizontal transmission (*β*), 2) the strength of density dependent mortality acting on juveniles (*k_J_*), 3) the strength of density dependent mortality acting on adults (*k_A_*), 4) the sensitivity of birth rates to past precipitation (*ρ*), and 5) the period of time over which past precipitation is averaged (*⍵*). In addition, testing against simulated data revealed that our method accurately estimates the time-averaged value of the composite epidemiological parameter *R*_0_, which measures the average number of new infections produced by a single infected individual (Figure 2). Details of simulation testing of our ABC methodology and calculation of *R*_0_ are described in the supporting online material.

**Figure 2.**
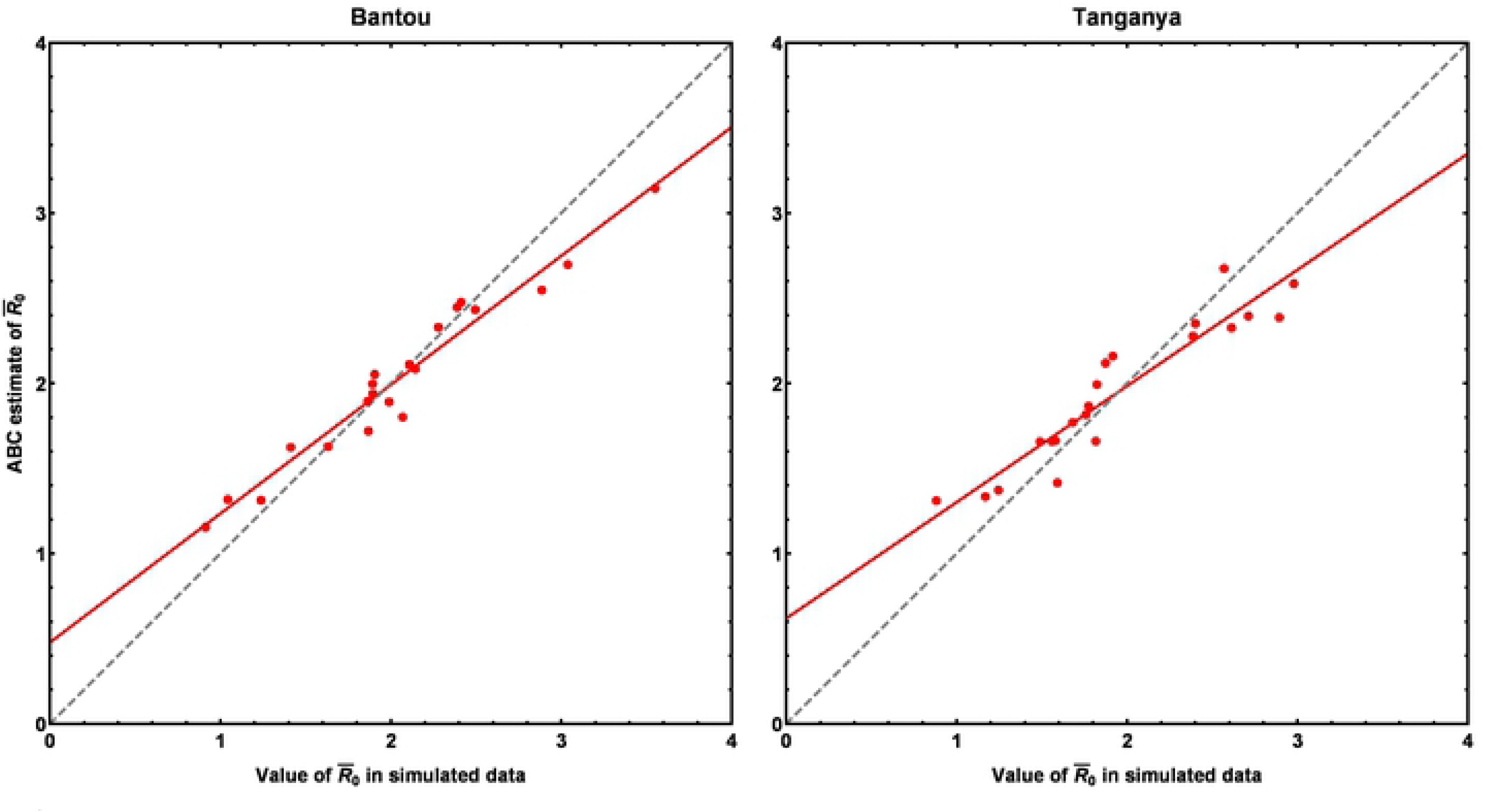
Comparison of time average values of *R*_0_ in simulated data sets (x axes) and the values of time averaged *R*_0_ estimated by our ABC approach (y-axes) when applied to the simulated data sets. The red dots indicate individual simulations, the red line the best fit to the dots, and the gray dashed line is the expected 1:1 relationship for a perfect fit. The equation for the line of best fit was given by *y* = 0.477 + 0.757*x* for Bantou and by *y* = 0.618 + 0.683*x* Tanganya where y is the value estimated by our ABC method and x is the true value in the simulated data set.

### Animal Studies

To make prior distributions for model parameters as accurate as possible, we supplemented information available from published studies with information derived from ongoing studies of a captive colony of *M. natalensis*. Specifically, unpublished data from a captive colony derived from Mali and housed at Rocky Mountain National Laboratories was used to refine prior distributions for maximum possible birth rate, *b_M_*, age at weaning, *α_PJ_*, age at first reproduction, *α_JA_* Similarly, the posterior distributions for the, and duration of viral shedding, *γ*.

#### Mastomys

All mice were bred and maintained under pathogen-free conditions at an American Association for the Accreditation of Laboratory Animal Care accredited animal facility at the NIAID and housed in accordance with the procedures outlined in the Guide for the Care and Use of Laboratory Animals under an animal study proposal approved by the NIAID Animal Care and Use Committee.

#### Ethics statement

*In vivo* studies were approved by the Institutional Animal Care and Use Committee of the Rocky Mountain Laboratories (RML). Animal work was conducted adhering to the institution’s guidelines for animal use, and followed the guidelines and basic principles in the United States Public Health Service Policy on Humane Care and Use of Laboratory Animals, and the Guide for the Care and Use of Laboratory Animals by certified staff in an Association for Assessment and Accreditation of Laboratory Animal Care (AAALAC) International accredited facility.

#### Biosafety

All work with infectious LASV and potentially infectious materials derived from animals was conducted in a Biosafety Level 4 (BSL 4) laboratory in the Integrated Research Facility of the Rocky Mountain Laboratories (RML), National Institute of Allergy and Infectious Diseases (NIAID), National Institutes of Health (NIH). Sample inactivation and removal was performed according to standard operating protocols approved by the local Institutional Biosafety Committee.

### Simulating Reservoir Vaccination Campaigns

When applied to the time-series data on rodent captures from the villages of Bantou and Tanganya in Guinea, our ABC approach results in a multi-dimensional posterior distribution describing the probability of various parameter combinations. Drawing parameter combinations from this posterior probability distribution and simulating forward in time while implementing vaccination allowed us to generate probabilistic predictions for the impact of different vaccination regimes. Specifically, we considered vaccination campaigns that rely on distributing vaccine-laced baits into both Bantou and Tanganya villages. We assume bait is consumed randomly at a rate, *σ*, by juvenile and adult rodents, but that pups do not consume vaccine baits prior to weaning. Pups can, however, be immunized through maternal transfer of protective antibodies with probability *M*, if born to a vaccinated mother. For simplicity, we assume consumption of a vaccine bait results in immediate, lifelong, complete immunity to Lassa virus. Because we assume baits are consumed by rodents at random, our model captures the reality that some proportion of vaccine bait is wasted on animals that are already infected with Lassa virus or recovered from prior Lassa infection and thus already immune. Specifically, the rate at which susceptible rodents of age class *i* (juvenile or adult) consume vaccine bait and become immune is given by:

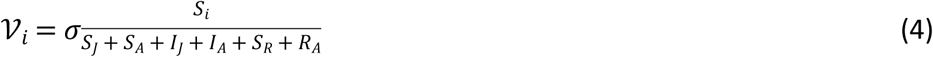

where the denominator is the total number of actively foraging rodents. The quantity 𝒱_*i*_ was capped at the number of susceptible individuals in class *i*.

We considered vaccination regimes that distributed between *σ* = 500-10,000 individual baits annually for an annual distribution period ranging from 7-168 days. In addition, we evaluated vaccination campaigns that began distributing bait in November (wet to dry transition) and those that began distributing bait in May (dry to wet transition). These temporal patterns were chosen because our ABC analysis revealed that reproduction in the *M. natalensis* population is maximal in November and minimal in May, and because previous work has demonstrated that the effectiveness of wildlife vaccination campaigns can be improved by focusing vaccination on periods of high reproduction (Schreiner et al. 2020). For each combination of bait number, baiting duration, and timing of bait distribution, we calculated the probability that Lassa virus was eliminated from both study villages (Bantou and Tanganya), and the average number of Lassa infected rodents over 100 replicate stochastic simulations.

## Results

### Parameter estimation for the Lassa virus pathosystem

We applied our ABC method to the rodent trapping data collected from the villages of Tanganya and Bantou in Guinea until the posterior distribution accumulated 28,042 points. Univariate analysis of marginal posterior distributions allowed us to calculate modal values and credible intervals for all model parameters (Table 3). Our analysis of simulated data revealed that only a subset of model parameters can be accurately and reliably estimated from the data, and accordingly, we focused on this subset of parameters here. Modal values for the remaining parameters differed little, if any, from the modal values of their prior distributions, consistent with our analyses of simulated data showing that little signal exists in the data for the value of these parameters (Supporting online material).

**Table 3.** Univariate modes and 95% credible intervals (highest posterior density) for model parameters

| Parameter | Mode | 95% Credible interval |
| --- | --- | --- |
| $b_M$ | 0.227 | {0.157, 0.284} |
| $\rho$ | 0.125 | {0, 4.50} |
| $\omega$ | 339.88 | {80.25, 365} |
| $k_{Bantou}$ | $1.676 \times 10^{-5}$ | $\{8.00 \times 10^{-6}, 2.82 \times 10^{-5}\}$ |
| $k_{Tanganya}$ | $2.08 \times 10^{-5}$ | $\{1.07 \times 10^{-5}, 3.37 \times 10^{-5}\}$ |
| $d_P$ | 0.0010 | {0.0006, 0.0015} |
| $d_J$ | 0.0033 | {0.0009, 0.0071} |
| $d_A$ | 0.0049 | {0.0004, 0.0104} |
| $\alpha_{PJ}$ | 0.0490 | {0.0317, 0.0701} |
| $\alpha_{JA}$ | 0.0100 | {0.0060, 0.0150} |
| $\beta$ | $1.56 \times 10^{-4}$ | $\{7.00 \times 10^{-5}, 2.54 \times 10^{-4}\}$ |
| $V$ | 0.0115 | {0, 0.161} |
| $M$ | 0.0105 | {0.0, 0.126} |
| $\gamma$ | 0.0450 | {0.0253, 0.0647} |
| $m_J$ | $3.28 \times 10^{-4}$ | {0, 0.004} |
| $m_A$ | $2.67 \times 10^{-4}$ | {0, 0.004} |
| $\tau_J$ | $4.08 \times 10^{-5}$ | $\{1.56 \times 10^{-5}, 8.04 \times 10^{-5}\}$ |
| $\tau_A$ | $5.09 \times 10^{-5}$ | $\{1.30 \times 10^{-5}, 9.70 \times 10^{-5}\}$ |

Posterior distributions for those parameters reliably estimable from the data were generally well-resolved. For instance, the rate of horizontal transmission of Lassa virus among juvenile and adult *M. natalensis* showed a well-defined peak at *β* = 1.56 × 10^− 4^ and the strength of density dependent mortality had clear univariate modes at *k* = 1.68 × 10^− 5^ in Bantou and *k* = 2.08 × 10^− 5^ in Tanganya (Figure 3). Similarly, the posterior distributions for the parameters *ρ* and *⍵* that quantify the sensitivity of *M. natalensis* birth rates to past precipitation were well-defined with modes at *ρ* = 0.125 and *⍵* = 339.88. Because these latter two parameters are functionally coupled with respect to their impact on the response of birth rates to seasonal fluctuations in precipitation (i.e., equation 2), however, we also explored their bivariate posterior distribution. Inspecting the bivariate posterior distribution yielded a slightly different mode for the parameter *⍵* (323.13) than when using its marginal distribution in isolation (Figure 4). To better understand the consequences of the full posterior distribution for likely patterns of seasonal reproduction in Bantou and Tanganya, we repeatedly sampled from the posterior distribution and calculated realized birth rates for Bantou and Tanganya villages over the period of the original field study using precipitation data for each village from the CHIRPS 2.0 database. This analysis demonstrated that the posterior distribution predicts modest levels of seasonal variation in birth rates, with peak and minimum reproduction in November and May, respectively (Figure 5).

**Figure 3.**
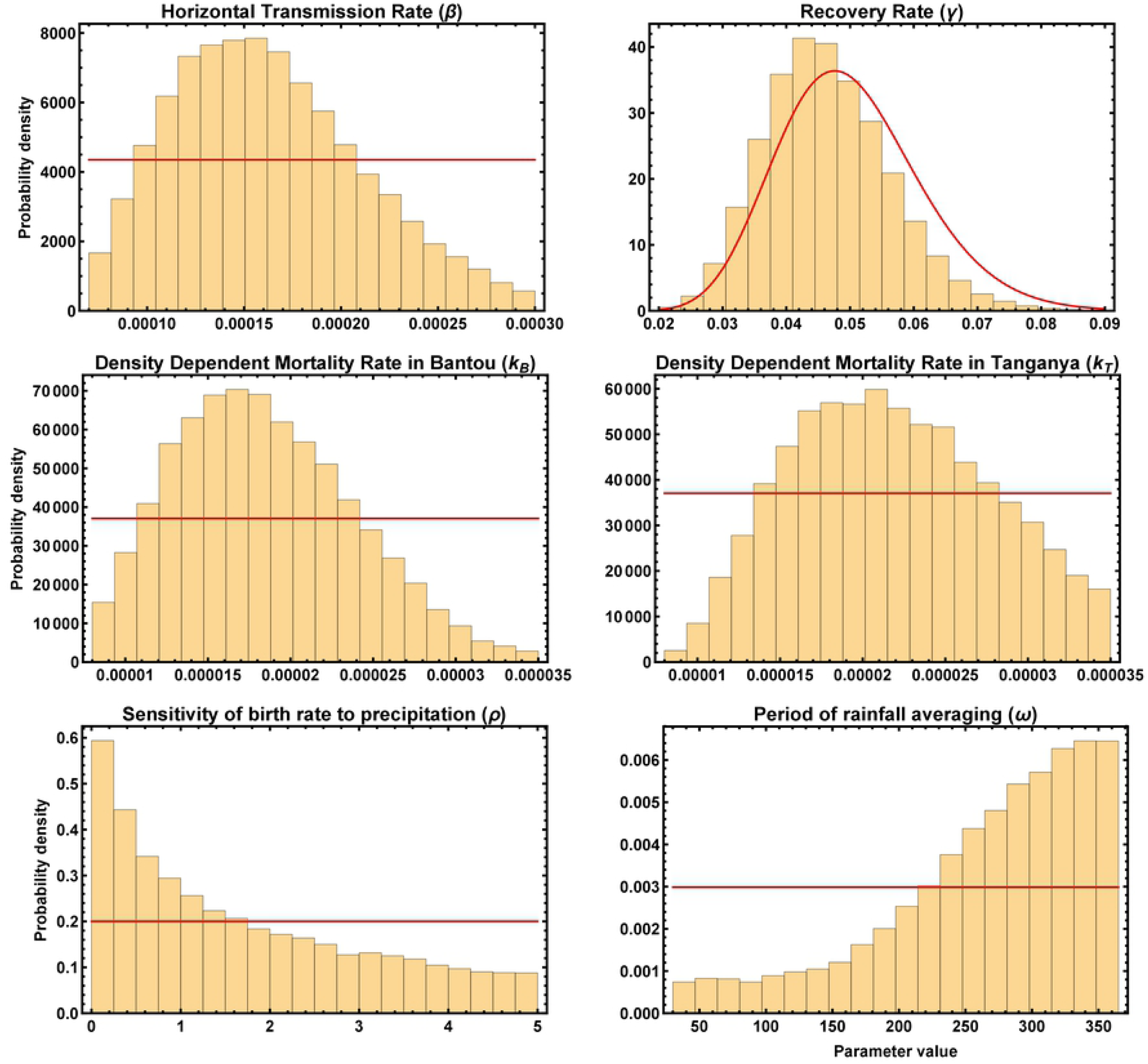
Posterior distributions for key model parameters estimated by our ABC method when applied to time series data from the villages of Bantou and Tanganya. The red lines show the prior distribution for each parameter and the gold bars the posterior probability density.

**Figure 4.**
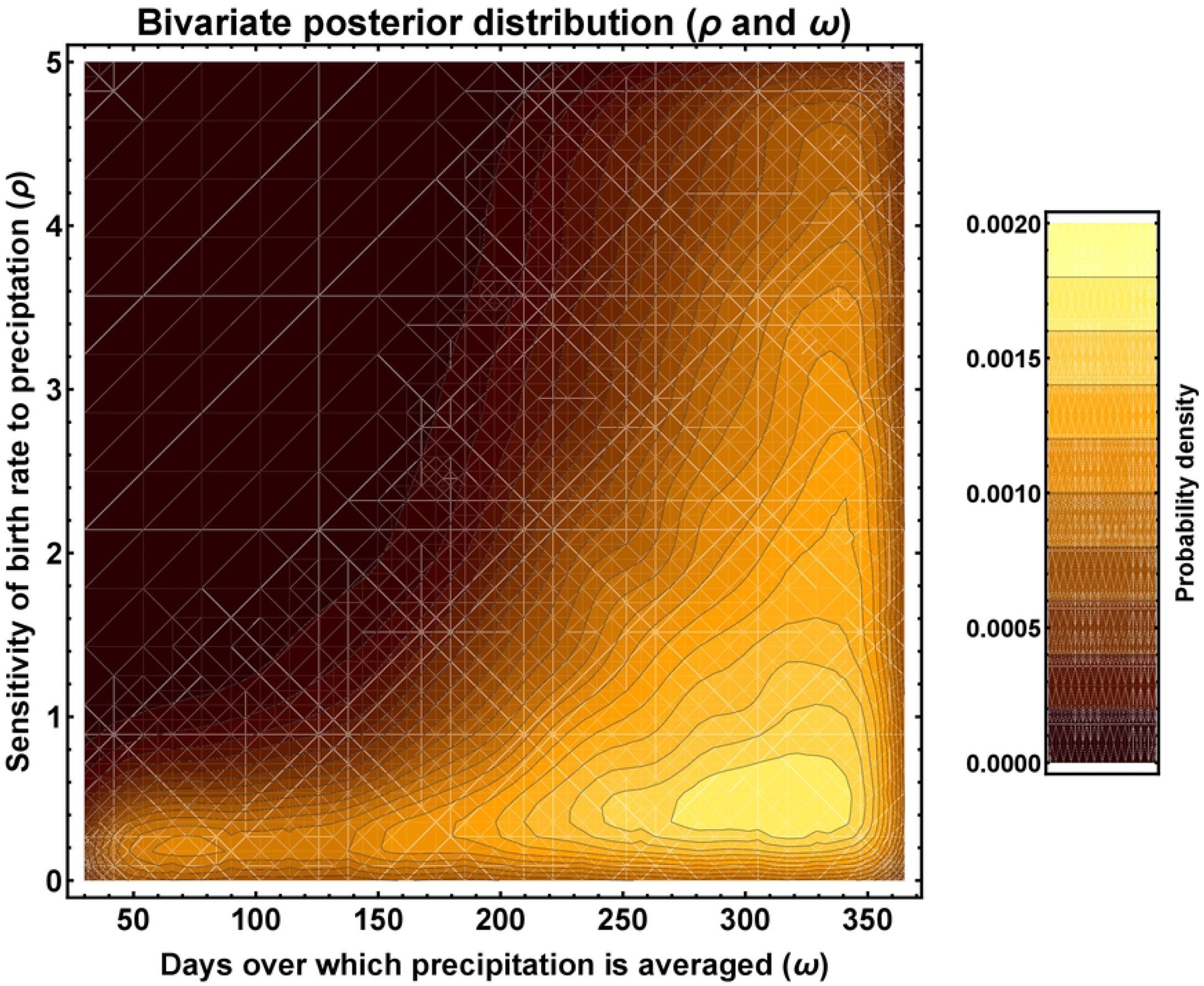
The bivariate posterior distribution for the parameters *ρ* and *⍵* that define the sensitivity of birth rate to patterns of past precipitation.

**Figure 5.**
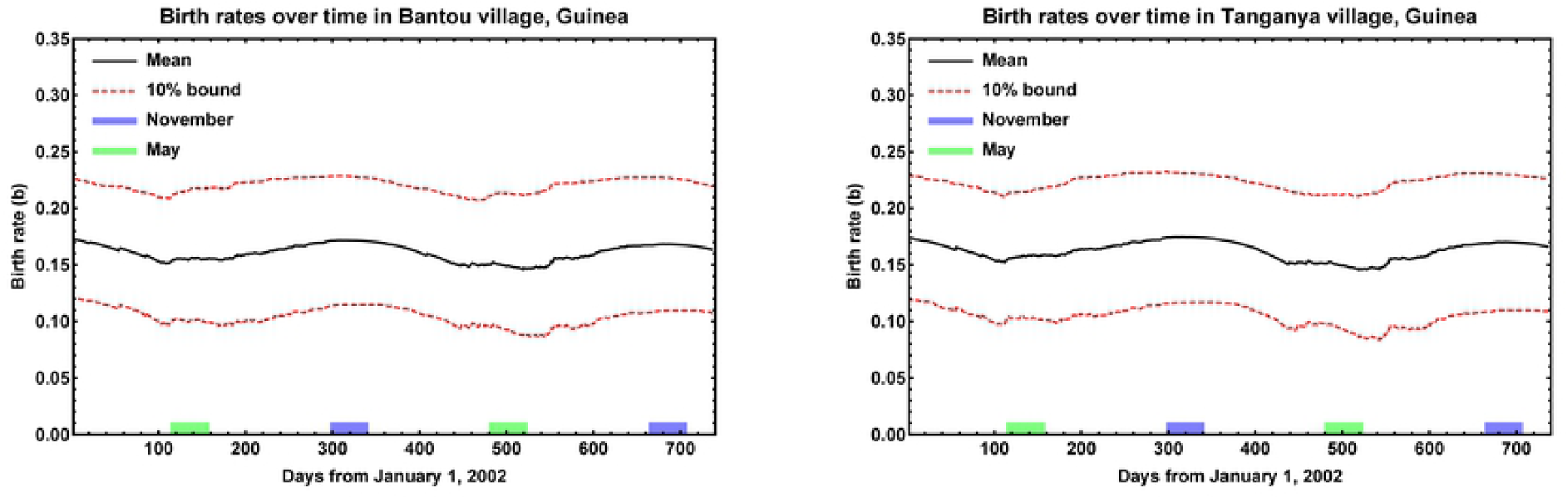
The predicted per capita birth rate of the *M. natalensis* population in the villages of Bantou and Tanganya over the period of the original field study. Average and prediction intervals were generated by: 1) drawing the parameters defining instantaneous per capita birth rates from the posterior distribution, 2) calculating the resulting per capita birth rate for the population over the period of the study using equation (2) and precipitation data for each village from the CHIRPS 2.0 database, 3) repeating this procedure for 1,000 random draws from the posterior distribution, 4) calculating the average birth rate at each point in time across the 1,000 replicates and 90% prediction interval.

To gain further insight into the epidemiological dynamics of Lassa virus within these villages, we generated posterior distributions for the time average value of *R*_0_ within each village (Figure 6). The modal values and credible intervals for *R*_0_ varied somewhat across villages, with modal values of 1.785 and 1.475 and credible intervals of {1.223, 2.346} and {1.202, 2.075} in Bantou and Tanganya, respectively. Using a classical result from epidemiological theory (Keeling and Rohani 2007), these values of *R*_0_ suggest that vaccination thresholds of 44.0% in Bantou and 32.2% in Tanganya would be sufficient for local elimination of Lassa virus from the reservoir population. Although these values suggest the feasibility of rodent vaccination, they rest on several important assumptions, including being able to vaccinate individuals prior to exposure by Lassa virus and population dynamics that are at a steady state. In contrast, vaccinating individuals prior to Lassa exposure is challenging and *M. natalensis* population dynamics are unlikely to be at a steady state. To explore how these features of the system influence the vulnerability of Lassa virus to rodent vaccination campaigns, we simulated reservoir vaccination campaigns using our mathematical model parameterized with the multi-dimensional posterior distribution derived from the villages of Bantou and Tanganya.

**Figure 6.**
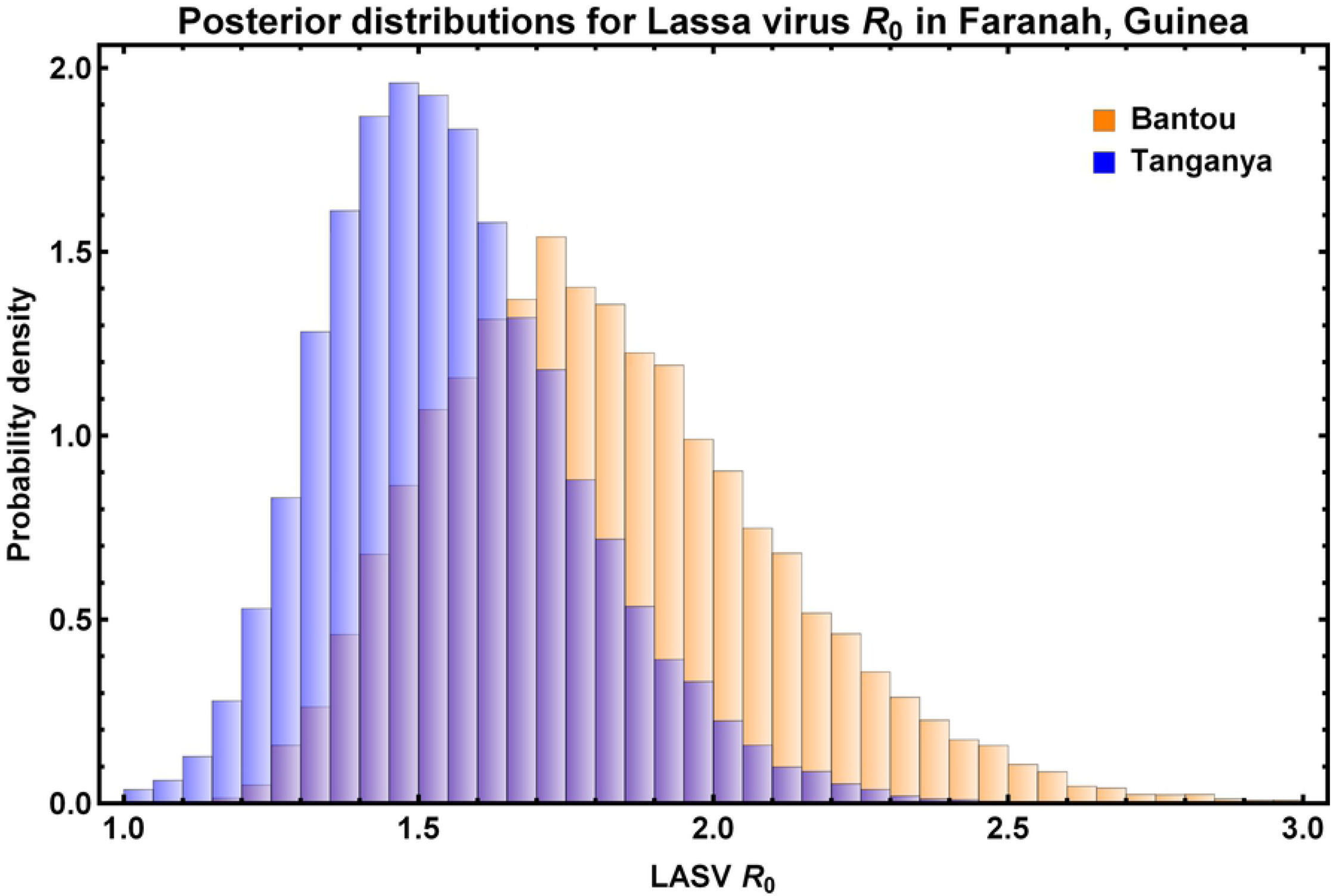
Posterior distributions for the time averaged value of *R*_0_ inferred by our ABC method for the villages of Bantou (orange) and Tanganya (blue). The modal values and credible intervals for *R*_0_ varied somewhat across villages, with modal values of 1.785 and 1.475 and credible intervals of {1.223, 2.346} and {1.202, 2.075} in Bantou and Tanganya, respectively.

### Simulating Reservoir Vaccination Campaigns

Results of our simulated vaccination experiments demonstrate that eliminating Lassa virus from both villages using distribution of conventional vaccine baits is extremely challenging and requires a level of bait distribution greatly in excess of existing wildlife vaccination programs. Specifically, our results show that the probability of eliminating Lassa virus from both Bantou and Tanganya villages is effectively zero when fewer than 4,000 baits are distributed in each village every year (Figure 7). Only when ≥ 4,000 baits are distributed annually over at least several weeks does the elimination of Lassa virus become a possibility (Figure 7). Results for reductions in the number of infected rodents are more encouraging, at least over the short term, with an appreciable (≈25%) reduction in the number of Lassa virus infected rodents achievable with distribution of ≤ 2000 baits per village and year (Figure 8). Notably, these reductions in the number of infected rodents were transient, with infections returning to near pre-vaccination campaign levels within 180 days following the end of the vaccination campaign (Figure 8, compare top and bottom rows). Comparing the results of vaccination campaigns that distribute baits when birth rates are at their peak (November) with vaccination campaigns that distribute baits when birth rates are at their minimum (May) revealed little, if any, impact of timing (Figures 7 and 8, compare left and right hand columns).

**Figure 7.**
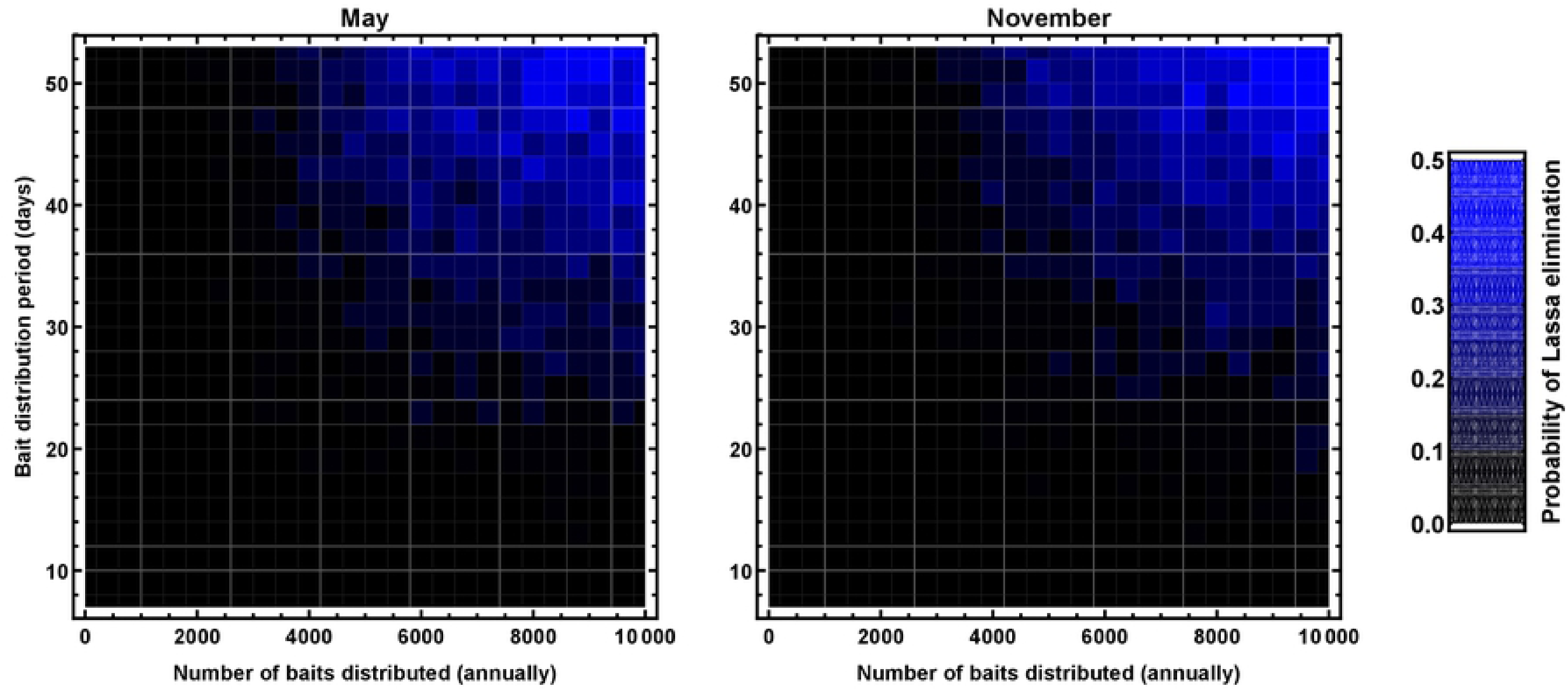
The proportion of simulated vaccination campaigns resulting in the simultaneous elimination of Lassa virus from the villages of Bantou and Tanganya as a function of the number of vaccine laced baits distributed per year (x axis) and the duration of bait distribution (y axis). The left hand column shows results for campaigns where vaccination occurs in November when birth rates are maximized (bait distribution begins November 1 of each year) and the right hand column campaigns where vaccination occurs in May when birth rates are minimized (bait distribution begins May 1 of each year).

**Figure 8.**
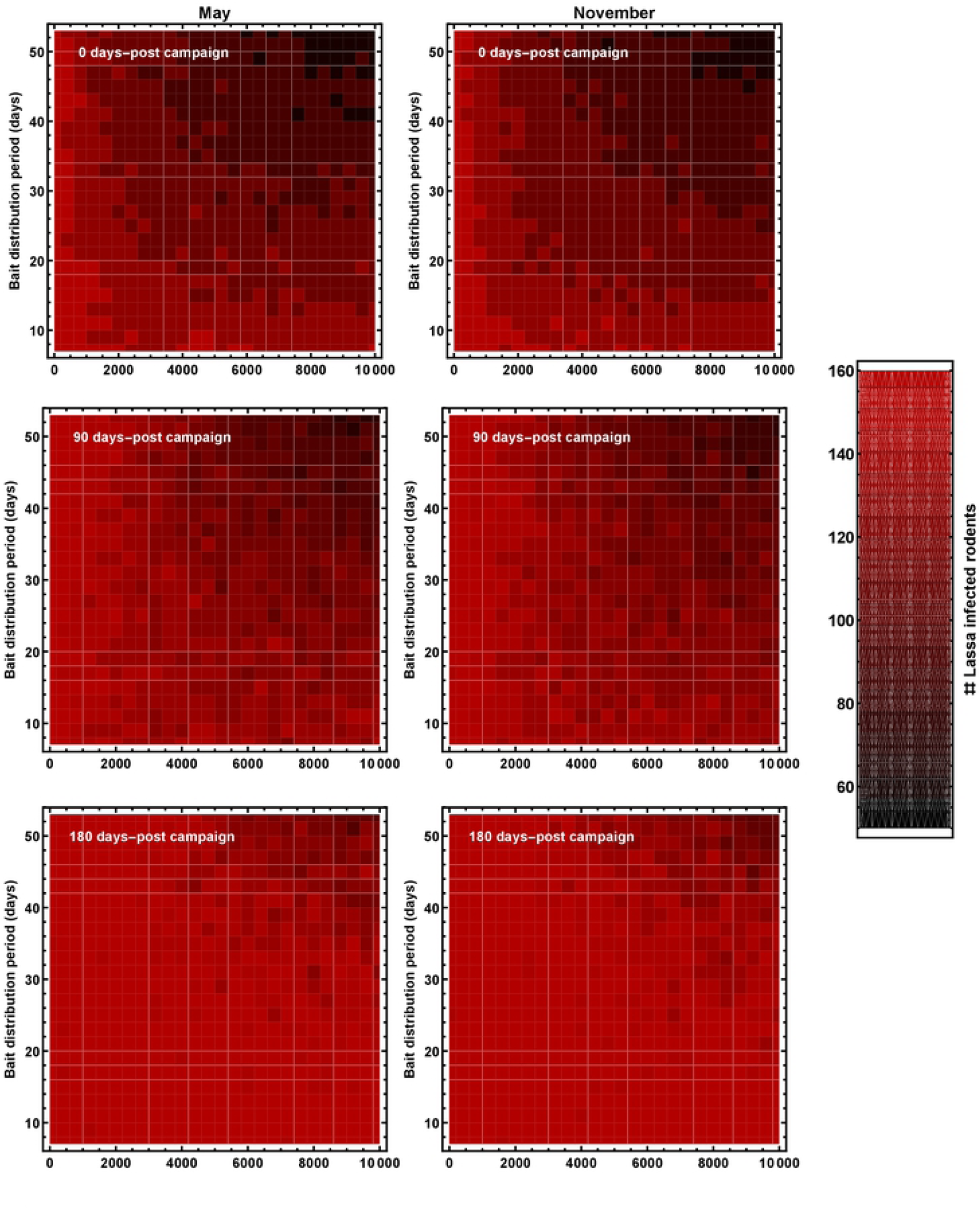
The number of *M. natanelsis* infected with Lassa virus 0, 90, and 180 days after the end of simulated vaccination campaigns (rows) as a function of the number of vaccine laced baits distributed per year (x axis) and the duration of bait distribution (y axis). The left hand column shows results for campaigns where vaccination occurs in November when birth rates are maximized (bait distribution begins November 1 of each year) and the right hand column campaigns where vaccination occurs in May when birth rates are minimized (bait distribution begins May 1 of each year). Any reductions in Lassa virus infection within the reservoir population accomplished by vaccination dissipate rapidly, with only modest reductions remaining 180 days after even the most intense vaccination campaigns cease.

## Discussion

We have developed an Approximate Bayesian Computation (ABC) approach for estimating demographic and epidemiological parameters of Lassa virus and its natural reservoir *M. natalensis* using time-series data on rodent captures. Extensive testing of our approach using simulated data sets demonstrates accurate estimation of key parameters such as the rate of horizontal transmission, the strength of density dependent mortality, and the sensitivity of birth rates to seasonal patterns of precipitation. In addition, our approach allows accurate estimation of the time averaged value of the composite parameter *R*_0_, quantifying the average number of new Lassa virus infections produced by a Lassa infected rodent introduced into an entirely susceptible population. Applying our approach to previously published data collected from two villages in Guinea (Fichet-Calvet et al. 2007; Fichet-Calvet et al. 2014) allowed us to develop estimates for epidemiological and demographic parameters, including robust estimates for the *R*_0_ of Lassa virus within its animal reservoir (*R*_0_ = 1.785 in Bantou and *R*_0_ = 1.474 in Tanganya). Although these estimates of *R*_0_ suggest Lassa virus may be vulnerable to vaccination campaigns targeting the rodent reservoir, *M. natalensis*, extensive simulated vaccination campaigns suggest distribution of conventional vaccine as bait may be ineffective unless an extremely high number of baits were regularly distributed.

There are at least two reasons reservoir vaccination is ineffective in our simulated vaccination campaigns. First, relatively large proportions of *M. natalensis* individuals within Guinea are known to be infected with Lassa virus or previously infected and recovered (9.1%-66.7% in the villages of Bantou and Tanganya). As a consequence, a large fraction of vaccine-laced baits will be consumed by rodents that are already infected by, or immune to, Lassa virus. Thus, to the extent that immunity to Lassa virus is lifelong as we have assumed here, only a relatively small fraction of distributed vaccine finds its way to its intended target. This contrasts with the case of rabies where relatively few animals are actively infected and high virulence prevents previously infected and immune animals from accumulating within the population (Hill et al. 1992). Second, the extremely high birth rate of the *M. natalensis* reservoir population causes immunity to wash out of the population very rapidly, with our simulations demonstrating that the impact of vaccine distribution generally lasts for less than 120 days once vaccine distribution ceases. Recent work using a different modeling framework and alternative assumptions came to a slightly more optimistic conclusion (Marien et al. 2019), most likely because direct vaccination of only susceptible individuals was assumed rather than the more realistic model of random bait consumption considered here.

There are at least two promising alternatives to the random distribution of vaccine laced baits: 1) coupling rodent removal through poisoning or trapping with distribution of conventional vaccine laced baits, or 2) vaccination through the use of scalable transmissible vaccines. In the first, the impact of rodent vaccination may be enhanced by coupling the distribution of vaccine laced baits with intensive rodent culling. By reducing the number of foraging rodents using relatively cheap culling methods (e.g., snap traps), this strategy may substantially reduce the number of vaccine baits required to reduce or eliminate pathogen infection, at least over the short term (e.g., Abdou et al. 2016). In some cases where culling has been employed and studied, however, it has been shown to have potentially counter-intuitive impacts, potentially increasing the prevalence of the target pathogen (Streicker et al. 2012; Amman et al. 2014). A second alternative to conventional vaccination relies on the use of transmissible vaccines capable of transmitting infectiously within the target population (Murphy et al. 2016; Nuismer et al. 2016; Bull et al. 2018). Transmissible vaccines have been show to substantially improve the likelihood of eliminating infectious disease in theoretical studies, but have been explored empirically in only a small number of cases (Bárcena et al. 2000; Angulo and Barcena 2007; Murphy et al. 2016; Bull et al. 2018). A transmissible vaccine targeting Lassa virus could potentially overcome the significant hurdles confronting traditional vaccination in this system. For instance, using formula derived in (Nuismer et al. 2016) along with the estimates for the time-averaged value of *R*_0_ in the Guinean populations we have derived here, shows that even a weakly transmissible vaccine with an *R*_0_ = 1.0 could reduce the number of baits that must be distributed by 56% in Bantou and 68% in Tanganya. A more strongly transmissible vaccine with an *R*_0_ in excess of 1.78 could allow Lassa virus to be autonomously eliminated from these populations.

Although our ABC approach allowed us to robustly estimate some epidemiological and demographic parameters, other parameters proved to be more challenging to estimate. Fortunately, most of these have been independently estimated in other studies (see Table 2), allowing their prior distributions to be defined with a relatively high degree of confidence. However, a number of parameters that our method is unable to estimate accurately have not been independently estimated (e.g., rates of Lassa virus vertical transmission) or have been estimated, but only in distant locations with substantial differences in rodent ecology (e.g., Tanzania)(Leirs et al. 1997; Van Hooft et al. 2008). Because of these limitations, there is some inevitable uncertainty in our estimates for model parameters. By generating our predictions for the efficacy of vaccination programs by repeatedly drawing parameters at random from the posterior distribution, however, this uncertainty is faithfully represented in predicted outcomes. In addition to assumptions about the likely values of a few poorly understood model parameters, our approach relies on a relatively simple mathematical model that simplifies population age structure, assumes density dependent infection and spatially homogenous transmission rates. Relaxing these assumptions could, in principle, alter our quantitative estimates for some model parameters (Borremans et al. 2017). Finally, our estimates of parameters are based on data collected over 14 years ago, creating the possibility that ecological, sociological, or evolutionary change in Lassa virus, the reservoir *M. natalensis*, or human populations with which the reservoir is commensal have caused values of important parameters to change.

Lassa virus is among the highest priority zoonotic pathogens identified by the World Health Organization and is a key emerging threat to human health (WHO 2018). Curtailing this significant threat to public health will likely require a combination of synergistic efforts including reservoir vaccination, targeted rodent control, reducing risky human behavior, and human vaccination campaigns. The ABC approach we have developed here provides a robust method for estimating key epidemiological parameters of Lassa virus and for predicting the likely effectiveness of these various types of intervention both individually and in combination. As additional field and experimental studies accrue, our ABC approach can be used to update and refine parameter estimates and predictions for intervention impacts. Only through rigorous evidence-based analyses and investigations of the impacts of all potential control options can global resources be effectively leveraged to combat this and other high-consequence threats to global health security.

## Acknowledgements

We thank James Bull for helpful comments on this work. The content of this publication does not necessarily reflect the views or policies of the Department of Health and Human Services, nor does the mention of trade names, commercial products, or organizations imply endorsement by the U.S. Government.

## Supporting Online Information

### Part 1: Evaluating performance of ABC approach using simulated data

Before applying our ABC method to real data, we evaluated its performance by testing against simulated data sets. Simulated data sets for the two Guinean villages were generated by: 1) drawing parameters at random from the prior distributions shown in Table 2 of the main text, 2) simulating population and epidemiological dynamics in these populations using the Gillespie algorithm and equations (1-2) with local precipitation data for each village from the CHIRPS 2.0 database, and 3) conducting simulated rodent trapping experiments. Simulated rodent trapping was conducted on the same dates as in the real dataset and the number of trap nights in each session was drawn at random from a uniform distribution on [800, 1200] to bracket values in the real data. The trapping rate parameter *τ* was set to 3.5 × 10^− 5^ for simulating data sets, a rate that yields biologically plausible values for true population density of M. natalensis within the Guinean villages that range between ≈ 600 ‒ 2,100 animals per village. In total, we generated 30 simulated data sets, each of which was then subjected to analysis using our ABC methodology.

To maximize the number of synthetic data sets we could analyze, we applied our ABC algorithm to each synthetic data set until 1000 points were accumulated in the posterior distribution. Of the 30 simulated data sets we analyzed, nine were terminated after failing to identify at least 50 hits within a week of computation. We continued analysis of the remaining 21 simulated data sets until at least 1000 points were accumulated in the posterior distribution. The small size of the posterior distribution used in our simulation testing should yield conservative estimates for how well our methodology is likely to work when applied to real data and run until a much larger number of points are accumulated in the posterior distribution. Once the ABC algorithm had been applied to each synthetic data set, we calculated the mode of the marginal posterior distribution for each model parameter and for the composite parameter *R*_0_.

Comparison of parameter values in the simulated data and the values inferred by our ABC methodology showed that five model parameters can be reliably estimated from the data: 1) the rate of horizontal transmission (*β*), 2) the strength of density dependent mortality acting on juveniles (*k_J_*), 3) the strength of density dependent mortality acting on adults (*k_A_*), 4) the sensitivity of birth rates to past precipitation (*ρ*), and 5) the period of time over which past precipitation is averaged (*⍵*) (Figure S1). Three other parameters showed a weak positive association between their values in the simulated data and their values estimated by our ABC algorithm (Figure S1). Remaining parameters showed no meaningful association between true and estimated values (Figure S2)

### Part 2: Calculating time-averaged *R*_0_

We calculated the time-averaged value of *R*_0_ using the next generation method and the density of susceptible individuals at each time point in the simulation (Diekmann et al. 1990). To calculate the *R*_0_ of system (1) at a single geographic location using the next-generation method, we rewrite equations 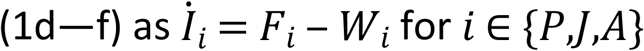 where *F_i_* gives the rate of new infections and *W_i_* gives the rate of change of *I_i_* by means other than new infections for age class *i*. This yields the following equations for *F_i_* and *W_i_*:

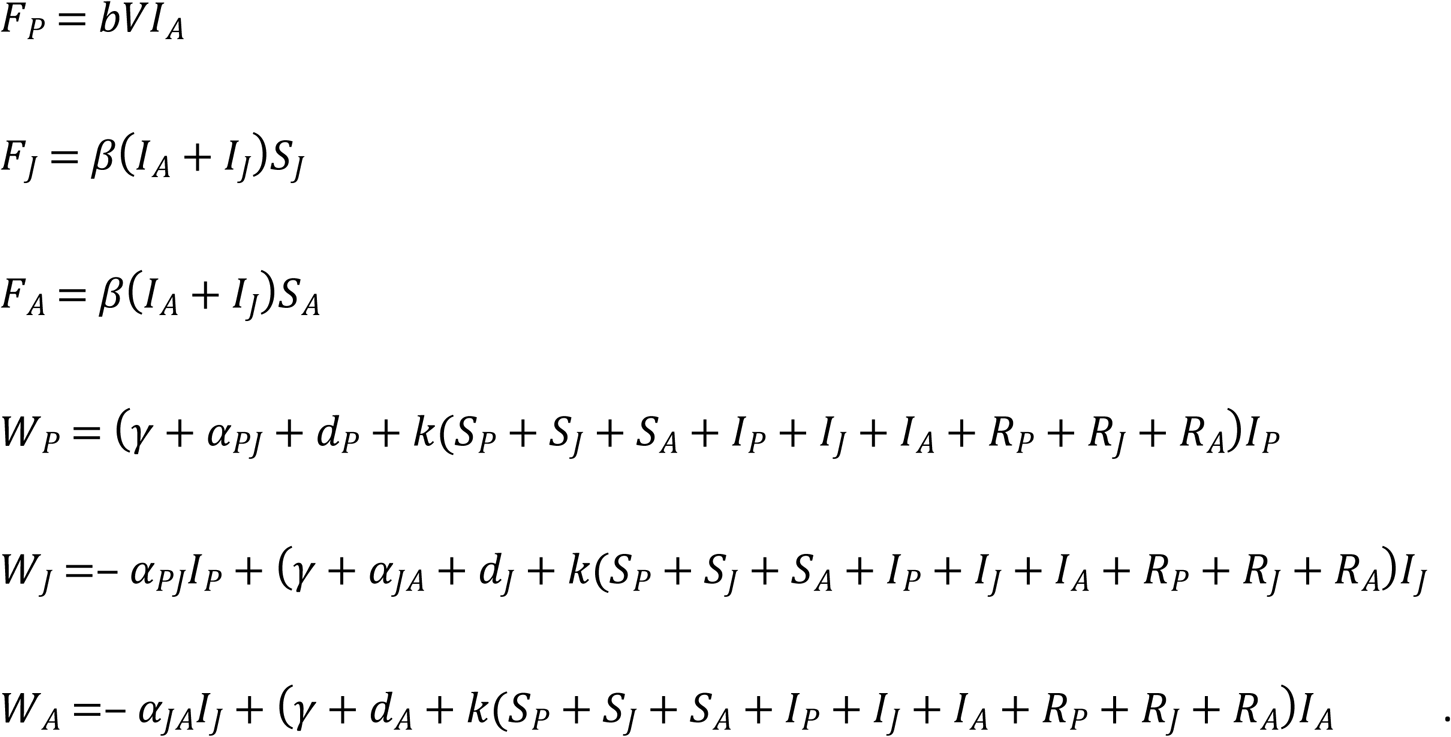

Let *N*_0_ = (*N_P_*,*N_J_*,*N_A_*)^*T*^ be the disease free equilibrium, in which all individuals are susceptible, and define the matrices 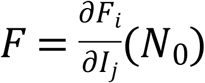 and 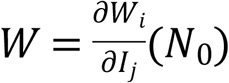 for *i,j* ∈ {*P*,*J*,*A*}. The next-generation matrix is given by *FW* ^‒ 1^. The spectral radius of *FW* ^‒ 1^ is the basic reproductive number:

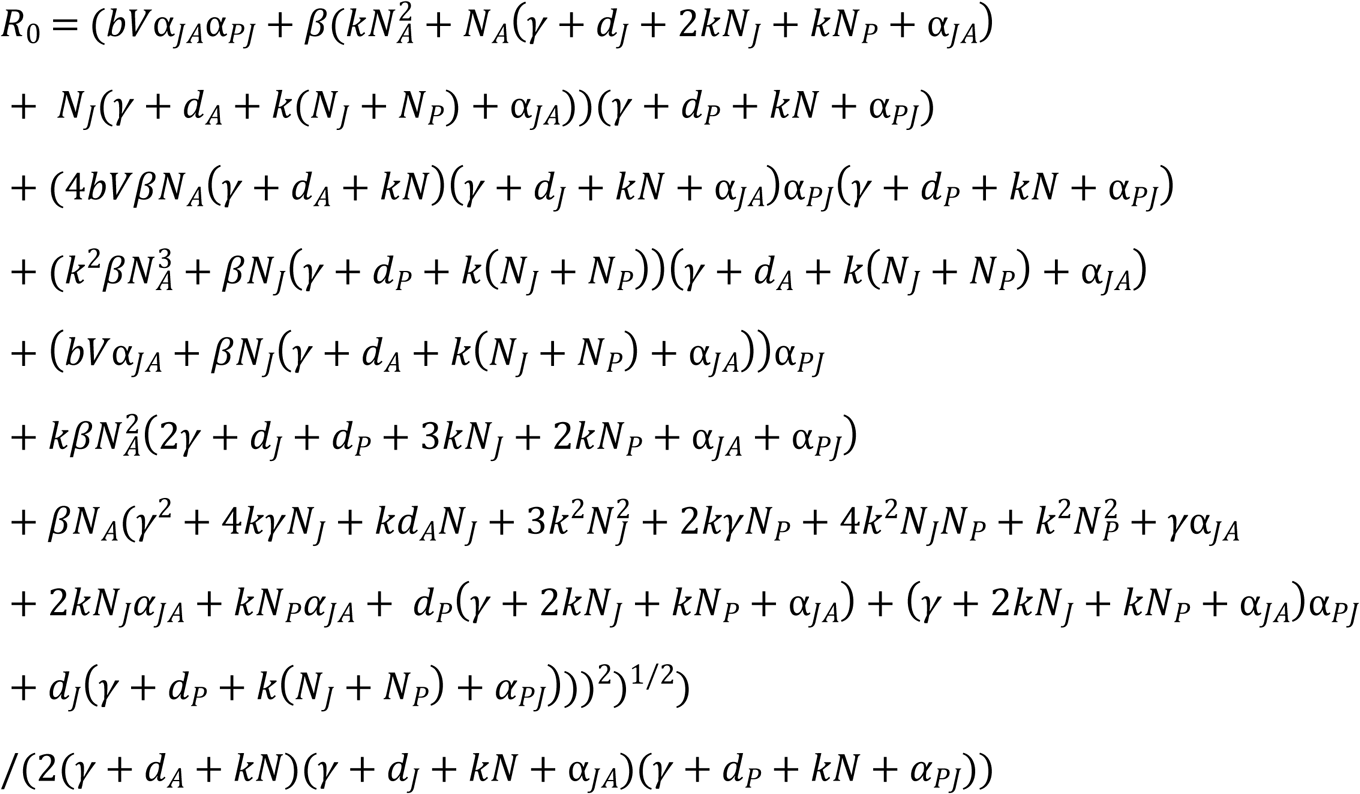

where *N* = *N_P_* + *N_J_* + *N_A_*.

An *R*_0_ value was calculated at each time point for each simulation in the posterior by plugging in the parameter values and the time-varying number of individuals in each age class *N_i_* for *i* ∈ {*P*,*J*,*A*}. The time-averaged *R*_0_ was then calculated by averaging these values for each simulation in the posterior.

---

**Figure S1.**
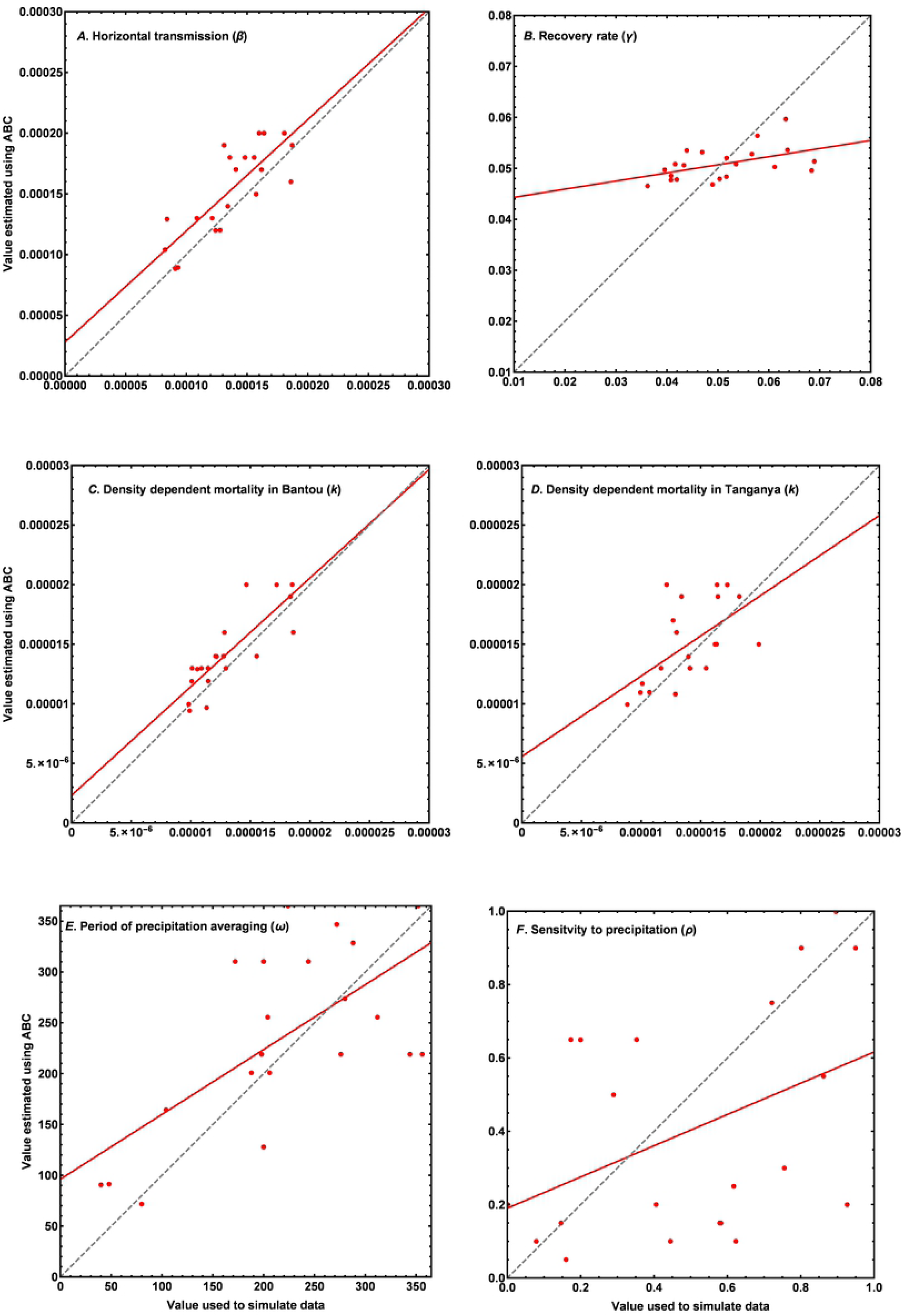

**Figure S2.**
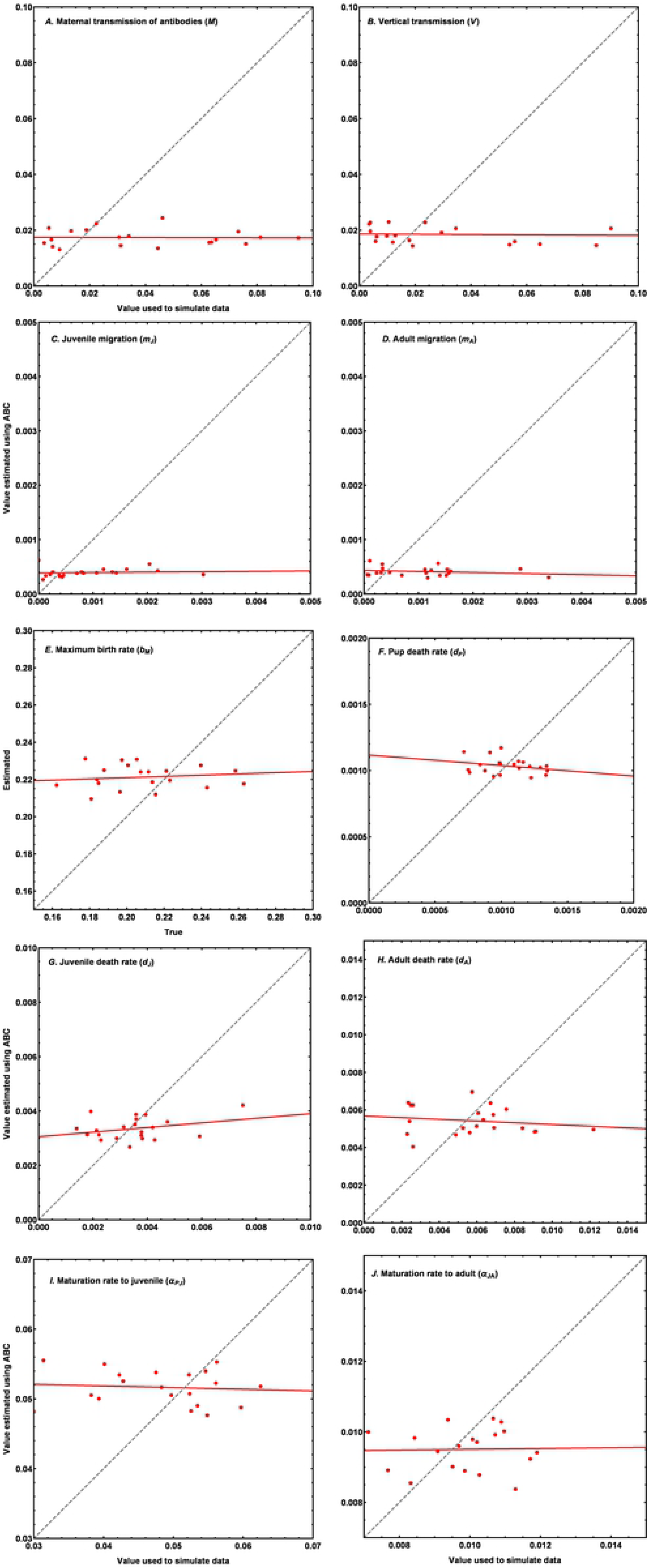

